# The effects of model complexity and size on metabolic flux distribution and control. Case study in *E. coli*

**DOI:** 10.1101/666859

**Authors:** Tuure Hameri, Georgios Fengos, Vassily Hatzimanikatis

## Abstract

Significant efforts have been made in building large-scale kinetic models of cellular metabolism in the past two decades. However, most kinetic models published to date, remain focused around central carbon pathways or are built around *ad hoc* reduced models without clear justification on their derivation and usage. Systematic algorithms exist for reducing genome-scale metabolic reconstructions to build thermodynamically feasible and consistently reduced stoichiometric models. It has not been studied previously how network complexity affects the Metabolic Sensitivity Coefficients (MSCs) of large-scale kinetic models build around consistently reduced models. We reduced the iJO1366 *Escherichia Coli* genome-scale metabolic reconstruction (GEM) systematically to build three stoichiometric models of variable size. Since the reduced models are expansions around the core subsystems for which the reduction was performed, the models are modular. We propose a method for scaling up the flux profile and the concentration vector reference steady-states from the smallest model to the larger ones, whilst preserving maximum equivalency. Populations of non-linear kinetic models, preserving similarity in kinetic parameters, were built around the reference steady-states and their MSCs were computed. The analysis of the populations of MSCs for the reduced models evidences that metabolic engineering strategies - independent of network complexity - can be designed using our proposed workflow. These findings suggest that we can successfully construct reduced kinetic models from a GEM, without losing information relevant to the scope of the study. Our proposed workflow can serve as an approach for testing the suitability of a model for answering certain study-specific questions.

**Author Summary:** Kinetic models of metabolism are very useful tools for metabolic engineering. However, they are generated *ad hoc* because, to our knowledge, there exists no standardized procedure for constructing kinetic models of metabolism. We sought to investigate systematically the effect of model complexity and size on sensitivity characteristics. Hence, we used the redGEM and the lumpGEM algorithms to build the backbone of three consistently and modularly reduced stoichiometric models from the iJO1366 genome-scale model for aerobically grown *E.coli*. These three models were of increasing complexity in terms of network topology and served as basis for building populations of kinetic models. We proposed for the first time a way for scaling up steady-states of the metabolic fluxes and the metabolite concentrations from one kinetic model to another and developed a workflow for fixing kinetic parameters between the models in order to preserve equivalency. We performed metabolic control analysis (MCA) around the populations of kinetic models and used their MCA control coefficients as measurable outputs to compare the three models. We demonstrated that we can systematically reduce genome-scale models to construct kinetic models of different complexity levels for a phenotype that, independent of network complexity, lead to mostly consistent MCA-based metabolic engineering conclusions.

## Introduction

Kinetic models of cellular metabolism can provide comprehensive understanding on the dynamics of the cell and its response to environmental changes and perturbations. In depth understanding of cellular metabolism can allow metabolic engineers to tailor cells according to sought specifications and objectives. This could enable the design of cell factories where flux is directed towards the production of biofuels, pharmaceuticals or other specialty chemicals. To be useful though, a kinetic model should represent the dynamics of the cell accurately enough to provide the required study-specific knowledge [1]. To date, important strides towards building large- and genome-scale kinetic models of metabolism have been made [2–5]. Despite the emergence of methodologies for building kinetic models, the research community knows that several challenges remain to be confronted.

With larger and better quality kinetic models, the mathematical representations become increasingly complex. Furthermore, the parameter sensitivities of systems biology models are in general “sloppy” [6]. Hatzimanikatis and coworkers noted that metabolic models are often built around certain central carbon pathways or, *ad-hoc* reduced models of genome-scale metabolic network models (GEMs) [7]. Such models do not account for the full information contained in the GEMs and, the *ad hoc* reduced models do not come with explicit explanations and justifications on how the model was reduced. Several studies have built kinetic models around *ad hoc* reduced models and computed Metabolic Sensitivity Coefficients (MSCs) for the system [1, 2, 8–10]. MSCs are desirable outputs of the kinetic models as they give insight into control patterns of the cell, assuming that the model is correct and accurate. However, Palsson and Lee showed that network complexity significantly affected the numerical values and the interpretation of MSCs [11]. Their study showed that three different red cell metabolic models produced MSCs that have opposite signs. This suggested that the analysis of incomplete metabolic models could lead to misleading and inaccurate information.

Palsson and Lee acknowledge that their models were reduced in an *ad hoc* manner to analyze how significantly network complexity can affect the MSCs of kinetic models [11]. However, nowadays algorithms for reducing GEMs are starting to emerge [7, 12, 13]. The NetworkReducer algorithm aims to reduce the network around certain “protected” metabolites and reactions by iteratively removing reactions that do not obstruct their activity [13]. Hatzimanikatis and coworkers developed the redGEM and lumpGEM algorithms which allow reduction of GEMs around selected subsystems by retaining linkages and the information captured in GEMs [7, 12]. The algorithm performs consistency checks with the GEM to ensure that the reduced model is consistent in terms of flux profiles, essential genes and reactions, thermodynamic feasible ranges of metabolite concentrations and ranges of Gibbs free energy of reactions. The redGEM and lumpGEM algorithms can be used to build thermodynamically feasible models with different levels of complexity consistent with the GEM for the same chosen subsystems. These algorithms open up the possibility to investigate how MSCs are affected by model complexity for consistently reduced models by building kinetic models around them.

We used the redGEM and lumpGEM algorithm to reduce the *E.coli* iJO1366 GEM to three different models, namely D1, D2 and D3, encompassing 271, 307 and 327 enzymatic reactions and 160, 188 and 197 metabolites, respectively. The thermodynamic formulation of the stoichiometric models allowed integration of fluxomics and metabolomics data for aerobically grown *E.coli* (S1 Table) [14]. Due to the topological differences between the three models, we proposed a technique for scaling up the flux profile and concentration vector reference steady-states from D1 into the larger models D2 and D3. This scale-up procedure ensures physiological equivalency of the models by assuring that their steady-states are numerically similar. All the three models satisfy thermodynamic constraints and are consistent with the GEM. We used the Optimization and Risk Analysis of Complex Living Entities (ORACLE) workflow to construct kinetic models for D1, D2 and D3 around their scaled reference steady-states. We fixed kinetic parameters from the smaller model into the larger one to further ensure equivalency of the models and hence a fair comparison. As integral part of the ORACLE workflow, we compute the MSCs for the stable kinetic models. We studied the MSCs across the three models and demonstrate that there is consistency amongst MSCs and that we can make metabolic engineering decisions, independent of model complexity.

## Results and Discussions

### Reduced *E.coli* models

We applied redGEM and lumpGEM algorithms [7, 12] to systematically derive modular, reduced, *E.coli* stoichiometric models (Methods) from the iJO1366 GEM [15]. We selected glycolysis, pentose phosphate pathway (PPP), tricarboxylic acid (TCA) cycle, glyoxylate cycle, pyruvate metabolism and electron transport chain (ETC) as the subsystems (as defined in the iJO1366 GEM [15]) around which reduction was performed to different degrees of connectivity, similarly to Ataman *et al*. [7]. These subsystems contain the 12 essential biomass precursors defined by Neidhart *et al* [16] and capture the central carbon metabolism of *E.coli*. Reduced stoichiometric models D1, D2 and D3 inter-connect the subsystems between each other with up to one, two and three reactions, respectively. Consequently, D1, D2 and D3 model cores generated via redGEM are constituted of 271, 307 and 327 enzymatic reactions and 160, 188 and 197 metabolites, respectively (Figure 1). The cores were connected to biomass production via lumped reactions, generated by the lumpGEM, to characterize the rest of the GEM (further discussion on lumped reactions around Figure 2 later in this section).

**Figure 1.**
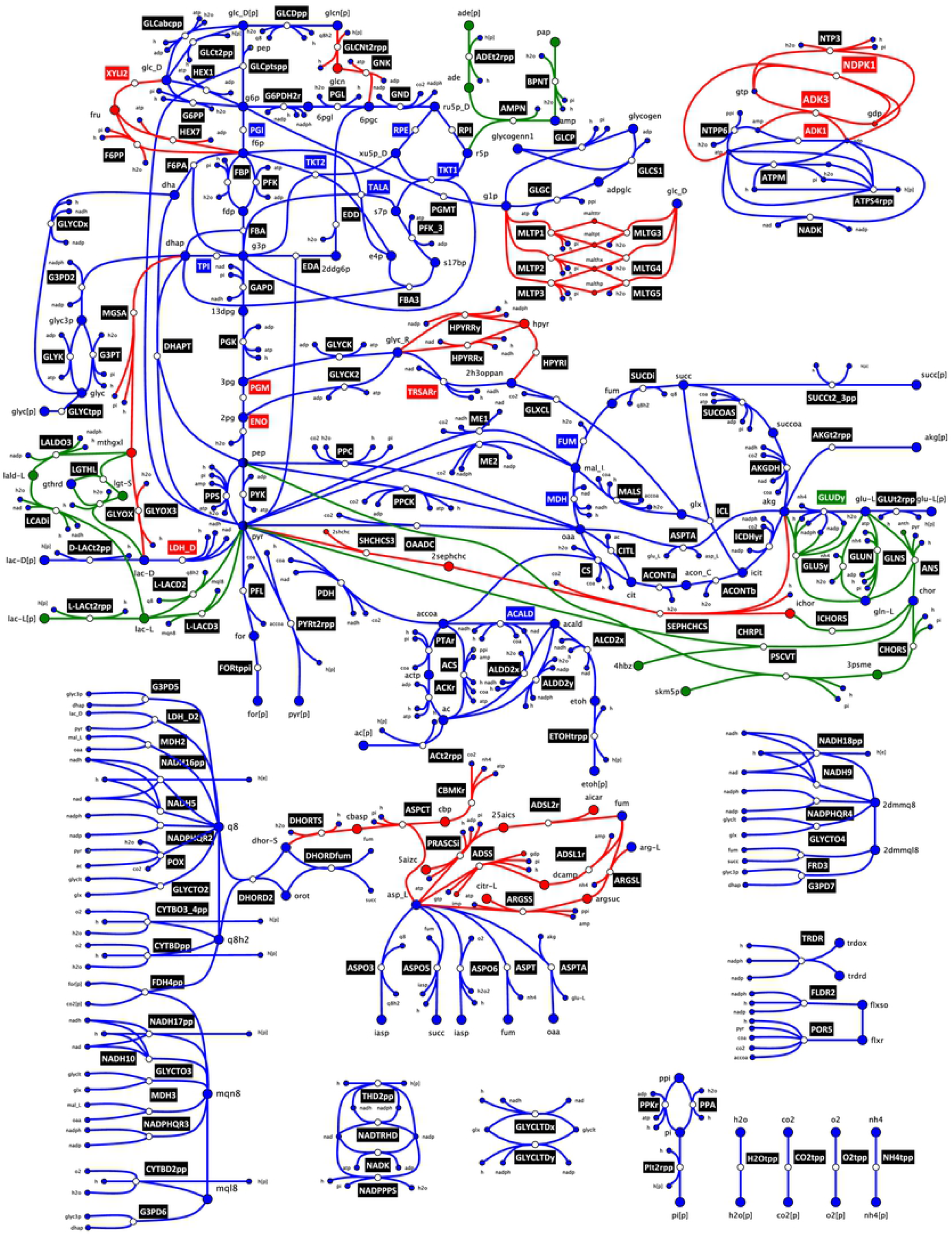
*E.coli* network diagram illustrating the core topologies studied for D1, D2 and D3. Edges represent the metabolic reactions and the nodes correspond to the metabolites. The reactions (edges) and metabolites (nodes) are coloured according to their pertinence to D1, D2 and D3, in blue, red and green, respectively. The reaction labels are coloured in black when the reaction is unidirectional. The bidirectional reactions’ labels are coloured if the reaction can operate in both forward and backward directions in D1, D2 or D3 in blue, red and green, respectively. The reactions that are bidirectional in a smaller model were also bidirectional in the larger model(s), as it is an expansion around the same core. Diagram does not include all the reactions of the systems.

The additional reactions in D2 include xylose isomerase (XYLI2), hexokinase D-fructose (HEX7) and D-fructose 6-phosphate phosphatase (F6PP), that connect D-glucose with D-fructose 6-phosphate via D-fructose. D2 also includes the maltodextrin system which connects the D-glucose to D-glucose 1-phophate via the maltodextrin phosphorylase and maltodextrin glucosidase reactions. In D2, dihydroxyacetone phosphate can react to methyglyoxal, which in turn can react to D-Lactate, providing increased connectivity between glycolysis and the pyruvate node. Additionally, pyruvate can react to 2-succinyl-5-enolpyruvyl-6-hydroxy-3-cyclohexene-1-carboxylate, which can react to form 2-oxoglutarate, thus connecting the TCA cycle with the pyruvate metabolism. D2 also includes three different ways to connect with two reactions from fumarate to L-aspartate, which – via argininosuccinate, adenylsuccinate and adenylosuccinate – further link the TCA cycle with the ETC. The adentylate kinase (ADK3), nucleoside-diphosphate kinase (NDPK1) and nucleoside-triphosphatase (NTP3) enzymes provide D2 model with additional flexibility in the system’s energy metabolism.

D3 has additional reactions enabling the transformation of methylglyoxal into D-lactate and L-lactate. Methylglyoxal is a hub metabolite that provides connectivity between upper glycolysis to the pyruvate node. The pyruvate and phosphoenolpyruvate nodes are connected to the TCA cycle via chorismate. Fruthermore, the glutamin and glutamate synthases provide additional flexibility in allowing conversion between L-glutamate and L-glutamine. In D3 the presence of AMP nucleosidase (AMPN) provides an additional connection between the PPP and the ETC. However, the expansion from D1 to D2 resulted in more central carbon metabolites that change in connectivity than the expansion from D2 to D3 (Figure 2). Several hub metabolites like methylglyoxal, isochorismate, pyruvate, D-lactate and 2-oxoglutarate change in connectivity between the three models.

**Figure 2.**
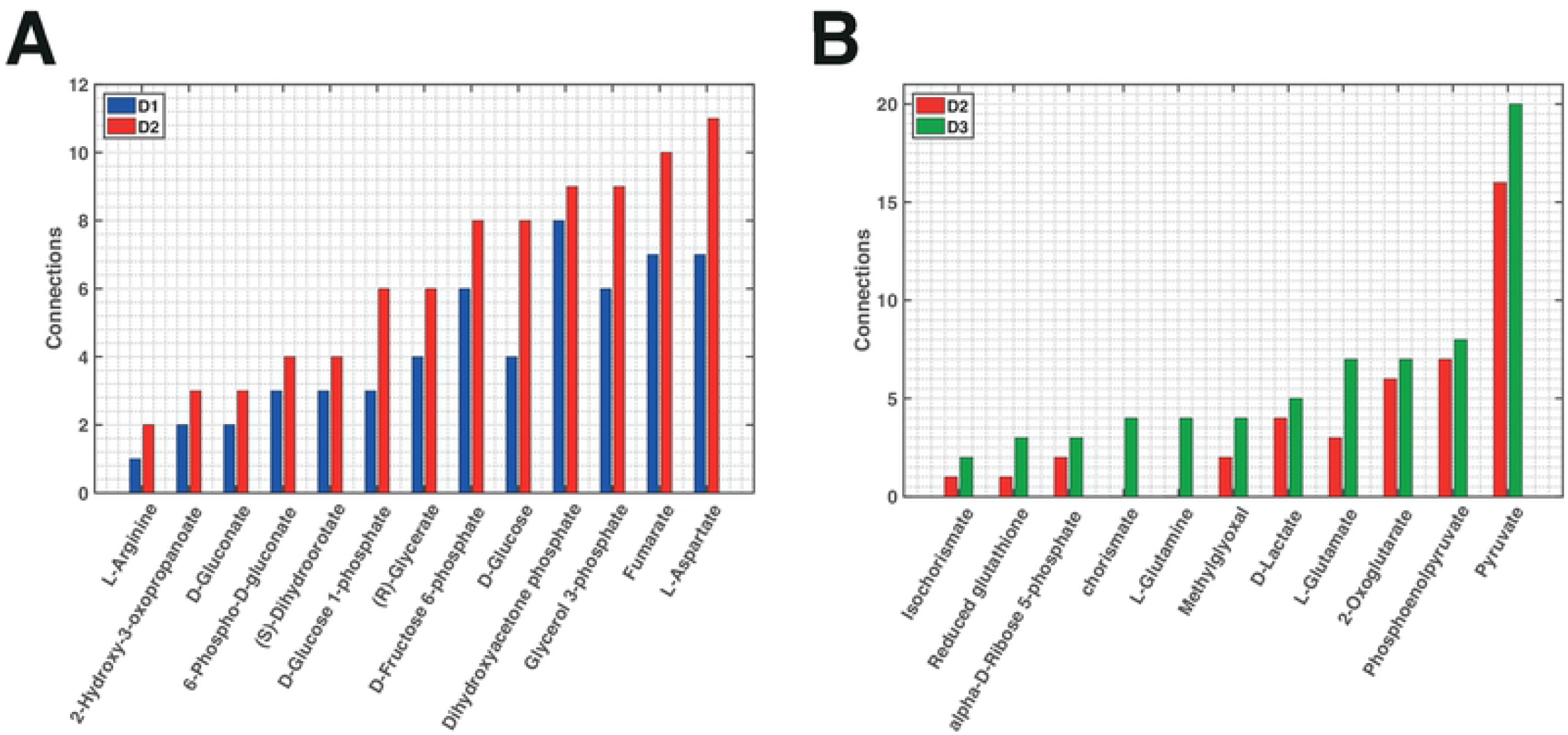
Metabolites connecting subsystems and the number of pairwise connections they achieve. Central carbon metabolites that change in connectivity (A) between D1 and D2, and (B) between D2 and D3.

### Thermodynamic-based variability analysis

Within the thermodynamic formulation [17] of the stoichiometric models D1, D2 and D3, we integrated fluxomics and metabolomics data for aerobically grown *E.coli* (S1 Table). Several assumptions were made on reaction directionalities, based on literature [14, 18–21], to further constrain the models (Methods). We performed a thermodynamic-based variability analysis (TVA) [22] on D1, D2 and D3 and we found they had 9, 17 and 18 bi-directional reactions, respectively (S2 Table).

Analysis of reaction flux ranges from TVA for D1, D2 and D3 revealed several considerable differences between the central carbon reactions. When comparing the ranges between D1 and D2, the largest differences were noted in adenylate kinase (ADK1), tartronate semialdehyde reductase (TRSARr), glucose-6-phosphate isomerase (PGI), phosphoglucomutase (PGMT), D-lactate dehydrogenase (LDH_D), triose-phosphate isomerase (TPI), phosphoglycerate kinase (PGK), glyceraldehyde-3-phosphate dehydrogenase (GAPD), phosphoglycerate mutase (PGM) and enolase (ENO) (S2 Table). Performing the same analysis between D2 and D3 reveals that only transketolase (TKT2) and glutamate dehydrogenase (GLUDy) changed considerably in ranges amongst central carbon reactions. In fact, GLUDy became bidirectional with this expansion. Other differences were noted in the peripheral reactions pertaining to additions from expansion from D2 to D3. Generally, most differences in the flux variability ranges resulted from the bypass routes that additional reactions provided, which resulted in certain reactions becoming bi-directional (Figure 1).

The TVA was also performed on concentration ranges of the models and we notice several differences in the allowable ranges of metabolite concentrations between D1, D2 and D3. Most noticeable concentration range differences between D1 and D2 occurred in D-glucose 6-phosphate, D-fructose 6-phosphate, fumarate, L-arinine and S-Dihydroorotate (S2 Table). However, between D2 and D3, noticeable differences were only noted in the ranges of D-glucose 6-phosphate and D-fructose 6-phosphate. The comparison of these TVA ranges in metabolite concentrations and reaction fluxes reveals that more considerable differences occurred between D1 and D2 than between D2 and D3.

### Model equivalency

Despite the inclusion of omics data for aerobically grown E.coli, D1, D2 and D3 remained underdetermined systems, resulting in the existence of multiple alternative --states that can characterize the studied E.coli physiology. A representative steady-state is required for the flux profile and for the metabolite concentration vector, to build a kinetic model around the selected steady-state. Furthermore, in light of benchmarking the outputs of kinetic models, the models are required to themselves be as close to each other as possible to allow for an unbiased comparison. Hence, their representative steady-states were kept similar so that the models describe the same operational state of the cell.

#### Scaling up steady-states

We sampled the flux and the concentration solution spaces for D1 and we used PCA to select representative steady-states (Methods). To preserve equivalency across the kinetic models, it was desirable that the flux profile and the concentration vector steady-states in D2 and D3 resemble the ones selected in D1. The topological modularity of the core models generated with redGEM eased the transferability of steady-states across models, allowing us to preserve similar values for fluxes and concentrations for the overlapping reactions of the three models.

We connected the core models to the biomass building blocks (BBBs), as defined by Neidhart *et al*. [16], via lumped reactions generated with the lumpGEM algorithm by applying approaches developed by Ataman and Hatzimanikatis [12]. A lumped reaction is a reaction that collapses a subnetwork of reactions into one mass-balanced reaction. D1-3 had 247, 189 and 196 lumped reactions, respectively. The models’ lumped reactions are indeed not the same across D1-3. Consequently, lumped reactions impose certain stoichiometric constraints that can require flux to pass through alternative metabolic routes within the models. For instance, a BBB can be produced by a completely different lumped reaction (Figure 3A), as we can generate it via a different subnetwork of reactions in the systems with larger cores. Thus, having distinct lumped reactions results in the redistribution of the flux profiles across models. An example of this is the hub metabolite methylglyoxal that provides new alternatives for lumped reactions in D2 and D3, thus contributing to differences in flux distribution across the models.

**Figure 3.**
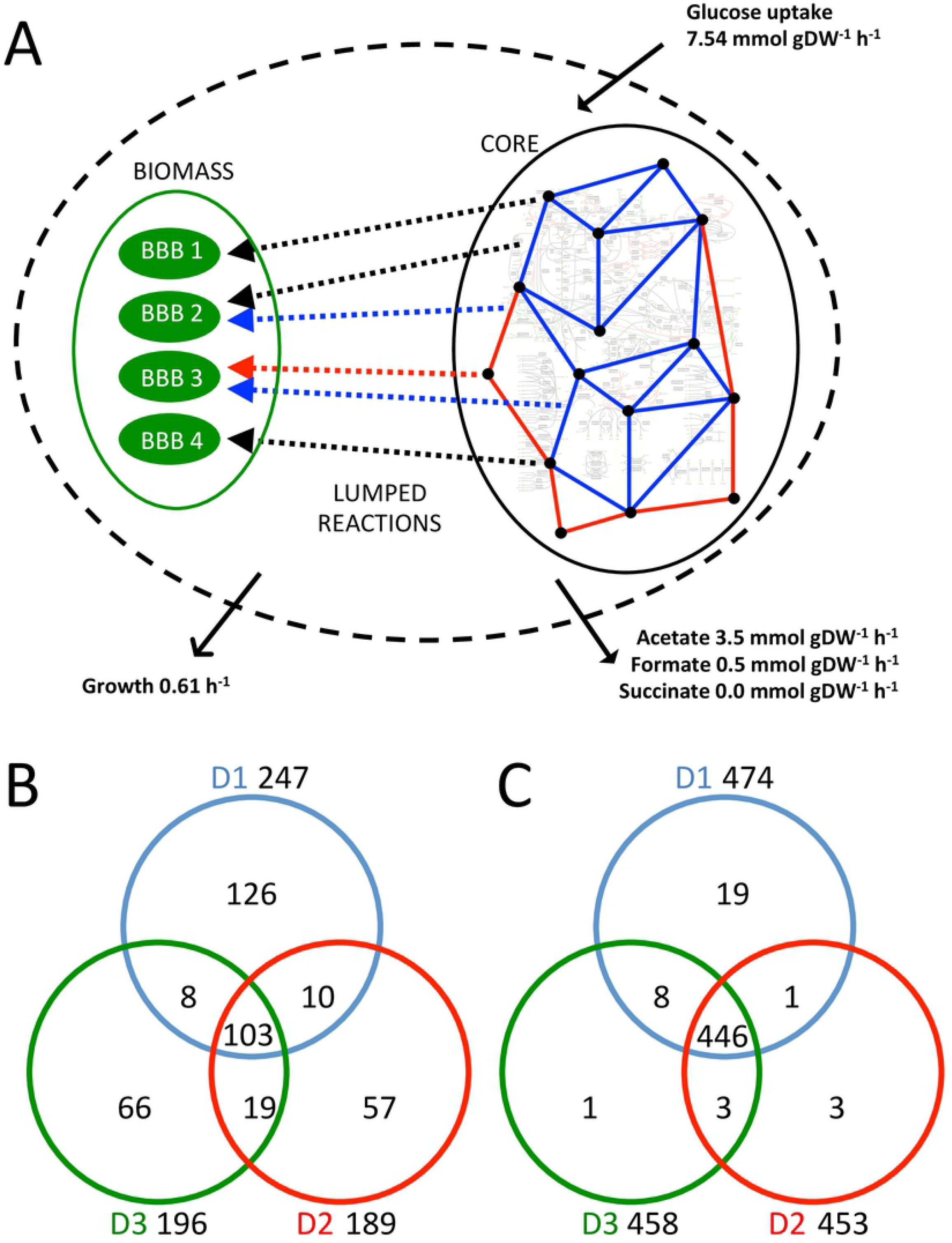
Illustration and analysis of core metabolism connections to biomass building blocks via lumped reactions. The schematic representation **(A)** of the studied metabolic networks where the edges are reactions and the nodes are metabolites. The blue edges and nodes belong to the core network of D1. The red edges and nodes correspond to additions from the D2 expansion. The dashed arrows are lumped reactions of the system, connecting the core to the biomass, where: black represents lumped reactions that are present in all the models, blue is for lumped reactions existing only prior to the expansion in D1 and red is for lumped reactions existing only after network expansion in D2. Biomass building blocks (BBBs) are represented by the green ovals. The black solid arrows indicate fluxomics data that were integrated for optimally grown *E.coli* [14]. Venn diagrams highlight differences in the lumped reactions of D1, D2 and D3 in terms of **(B)** subnetworks and in terms of **(C)** reactions.

We studied the lumped reactions and in D1-3 and observed that 103 were common between the models (Figure 3B). D1, D2 and D3 have 126, 57 and 66 lumped reactions that are unique to themselves. D1 requires considerably more lumped reactions in order to produce the BBBs from the core subsystems. If we consider the lumped reactions as subnetworks of reactions, 474, 453 and 458 reactions are used to build the lumped reactions of D1-3, respectively (Figure 3C). Interestingly, 446 reactions are common between the pools of reactions that constitute the lumped reactions of D1-3.

In order to ensure equivalency between D1-3, we proposed a procedure that uses a Mixed Integer Linear Programming (MILP) formulation that imposes similarity between the representative steady-states of the models (Methods). The D2 fluxes of central carbon reactions are within below one percent deviation from the reference flux of D1 (S3 Table). The only central carbon fluxes in D3 that deviate from D2 reference flux with more than one percentage are transaldolase (TALA) and xylose isomerase (XYLI2) with 4.5% and 33.2% respectively. The concentration profile of D2 is within one percent of D1 reference steady-state, except for ADP, CoA, S-dihydroorotate and L-glutamine with 16%, 45%, 303% and 94% deviations from D1. On the other hand, the D3 metabolite concentration steady-state is within one percentage from the D2 metabolite concentration vector. The modularity and the consistency of redGEM and lumpGEM algorithms in GEM reduction allowed the steady-states to be transferred and communicated between models efficiently.

#### Equivalence in kinetic parameters

We constructed kinetic models around the selected reference steady-states of D1-3 using the ORACLE workflow [3, 23–25]. Uniform Monte Carlo sampling of the degrees of saturation of the enzyme active sites allowed us to study the kinetic parameter space, as proposed by Hatzimanikatis and colleagues [23]. The local stability of the models generated was tested by verifying that the eigenvalues are not positive. We first sampled 50’000 stable kinetic models for D1. To ensure equivalency at kinetic parameter level between D1-3, we adapted the ORACLE workflow to allow fixing the sampled saturation states from one model to another (Methods). From the 50’000 stable D1 kinetic models, we found 96.1% (48’080) to be stable in D2, of which 98.4% (47’299) were stable in D3. We then computed the MSCs for these stable models in order to compare how MCA-based decisions are affected by metabolic network size.

### Consistency in MCA across models

#### Ranking enzymes for flux control

Some fundamental cellular tasks for a given physiology include metabolite excretion, substrate uptake and cellular growth, *μ*. As we studied the physiology of optimally grown E.coli, we considered control over *μ* across models to assess the consistency in conclusions based on MCA outputs. The flux control coefficients (FCCs) of *μ* were ranked for D1-3 based on their absolute means across stable models. The models were compared pairwise in increasing order of size (i.e. D1 versus D2, and D2 versus D3) to assess the impact of systematic network expansion on MCA (Figure 4).

**Figure 4.**
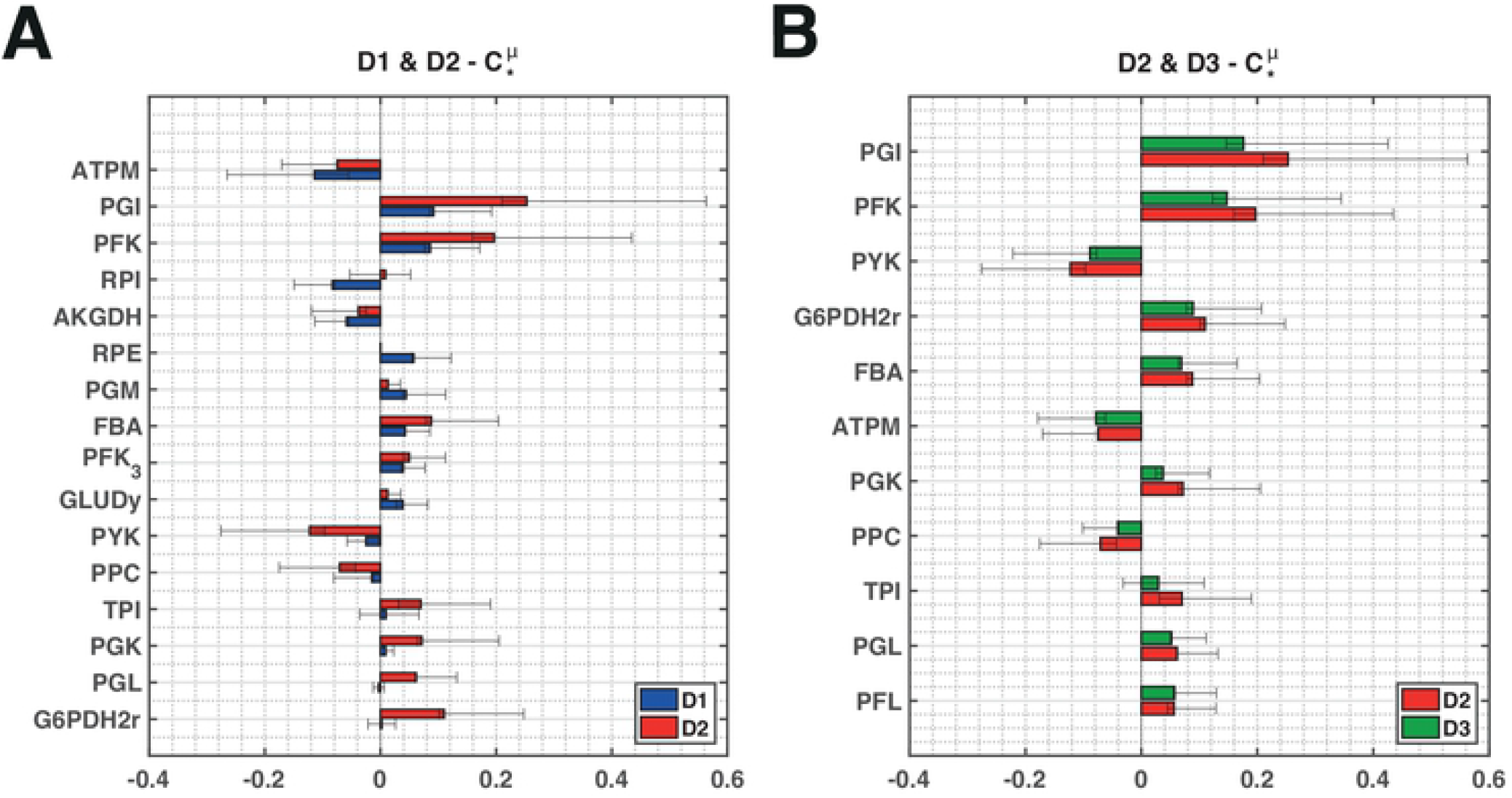
Cellular growth (*μ*) control across models. Pairwise illustration of the union of the top 7 enzymes across the models in terms of absolute control over cellular growth for **(A)** D1 versus D2, and **(B)** D2 versus D3. The whiskers give the upper and lower quartiles of the FCC populations and the bars give the means.

The cellular growth FCCs with respect to PGI, phosphofructokinase (PFK) and ATP maintenance (ATPM) are the most consistent in terms of sign and magnitude when comparing D1 with D2 (Figure 4A). Pyruvate kinase (PYK), fructose biphosphate aldolase (FBA) and 2-oxogluterate dehydrogenase (AKGDH) are also in agreement in terms of sign but magnitude can differ significantly. Some enzymes have control in D1 but no control in D2, such as ribulose 5-phosphate 3-epimerase (RPE) and phosphoglycerate mutase (PGM). Others, vice versa, have control in D2 but no control in D1 such as phosphoglycerate kinase (PGK) and glucose 6-phosphate dehydrogenase (G6PDH2r). Ribose-5-phosphate isomerase (RPI), on the other hand, has opposing control on cellular growth in the two models. Differences in FCCs of cellular growth between D1 and D2 suggest that the modular expansion of D1 to D2 significantly affects the control scheme.

We then compared top cellular growth FCCs in D2 and D3, which are in great sign and magnitude agreement (Figure 4B). PGI, PFK and PYK are the top three enzymes in terms of cellular growth control according to both D2 and D3. The consistency between these FCCs suggests that the modular expansion of D2 to D3 does not affect the control pattern as significantly as the network expansion from D1 to D2. An analogous analysis was carried out for the flux control of glucose uptake and several other cellular excretions (S4 Figure), and we observed a similar trend. The differences in control patterns appear to be significant when expanding from D1 to D2, but of lesser importance when expanding from D2 to D3. This finding could suggest that entire genome-scale kinetic models are not necessary to capture the essential physiological features of a cell as long as the model reduction is done systematically around carefully selected subsystems of study interest. Additionally, this could mean that D1 is possibly missing on some important information for performing MCA. Clearly, a study-specific resolution criterion in terms of model size/complexity that has to be met needs to be established before a model is used for further analysis.

#### MCA consistency across reduced models

As the study above revealed, certain flux control patterns can change significantly between models due to network complexity. We tried to locate, analyze and understand the differences and the similarities in MSCs that occur due to the topological alterations in kinetic model complexity. According to MCA theory, the FCCs conform with the summation theory [26, 27]. We proposed a deviation index (DI) that provides a quantitative measure on how much a reaction’s FCCs differ between two models, postulated from the summation theory (Methods). The DI served as a metric to classify reactions with respect to their consistency in FCCs across the reduced models.

We estimated the DI of 271 common enzymatic reactions when expanding from D1 to D2 to predict deviations in FCCs for the system. Reactions with the lowest DI (0-25 percentile) were mostly from the central carbon metabolism (Figure 5). The reactions with the highest DI (75-100 percentile) were mostly located in the ETC. The only central carbon metabolism reactions having a high DI were TALA, acetyl-CoA synthase (ACS), phosphoenolpyruvate synthase (PPS) and NAD malic enzyme (ME1). TALA produces D-fructose 6-phosphate and, PPS and ME1 involve transformation of pyruvate. D-fructose6-phosphate and pyruvate are both central carbon metabolites around which the expansion adds reactions (Figure 2 and 5). ACS is only one reaction away topologically from pyruvate, around which the expansion adds a reaction (Figure 2 and 5).

**Figure 5.**
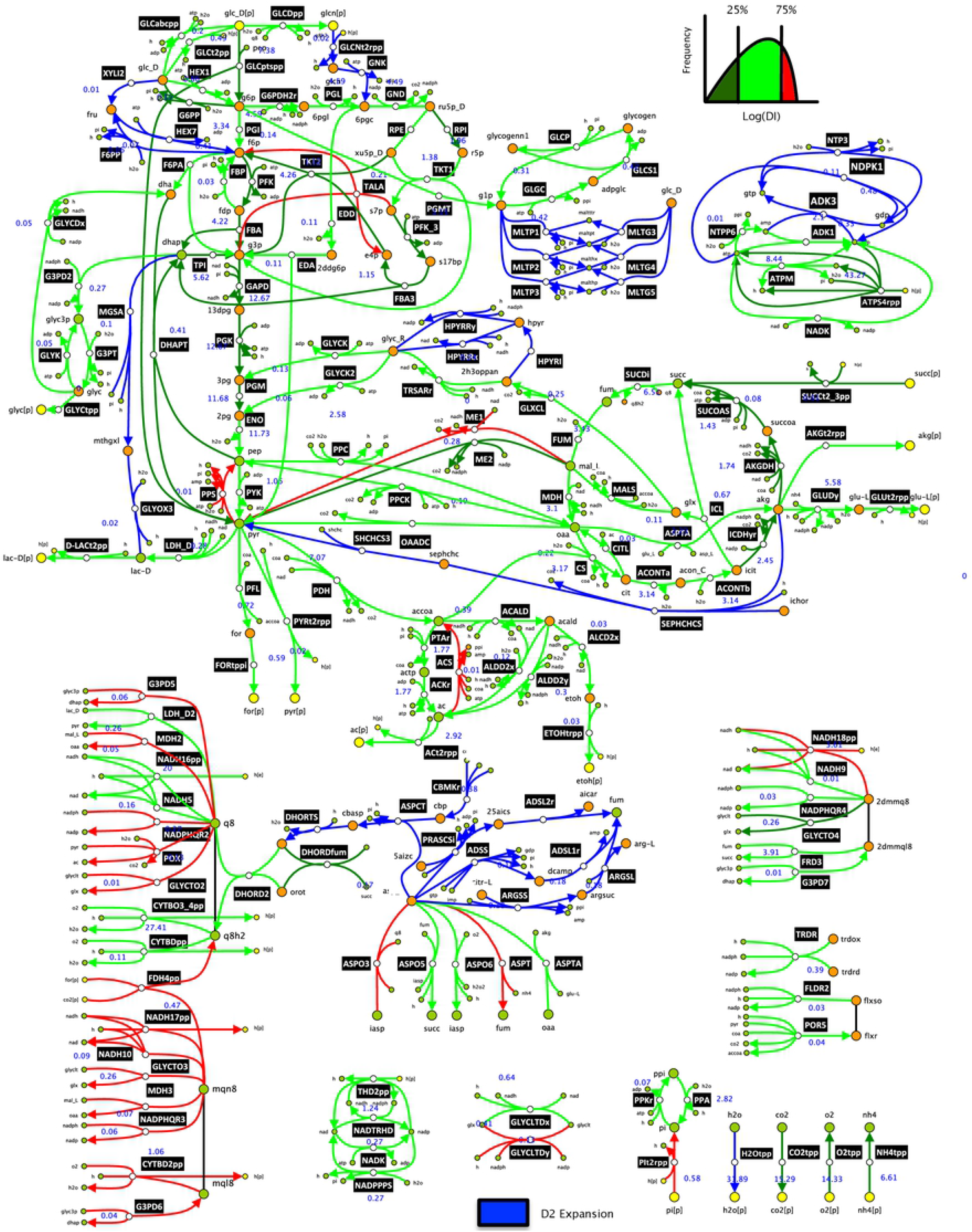
*E.coli* network diagram illustrating the deviation index (DI) of reactions when scaling up from D1 to D2. The flux values of the D2 flux profile (reference state) are given for the reactions in mmol/gDW/h next to their labels. The reactions added by the modular redGEM expansion from D1 to D2 are given in blue and the 271 enzymatic reactions in common between D1 and D2 are color-coded based on the DI. The reactions were categorized into low DI (0-25 percentile), medium DI (25-75 percentile) and high DI (75-100 percentile), and are given in dark green, light green and red, respectively. Diagram does not include all the reactions of the systems.

We repeated the above analysis for D2 and D3, where we analogously compute the DIs for the 307 common enzymatic reactions (Methods). Similar observations were made for the reactions having low DIs (0-25 percentile) as most were located in central carbon metabolism, within the subsystems around which reduction was performed (Figure 6). The reactions with higher DIs (75-100 percentile) are predominantly located around the ETC, with the exception of several reactions pertaining to central carbon metabolism. As with the previous analysis of D1 versus D2, ME1 and ACS had high DIs. The D3 expansion adds reactions around pyruvate (Figure 2 and 6), which could explain this observation. Aspartate transaminase (ASPTA), phosphoenolpyruvate carboxykinase (PPCK) and succinate dehydrogenase (SUCDi) from the central carbon metabolism exhibited high DIs (Figure 6). ASPTA is directly connected via 2-oxoglutarate with NADPH glutamate synthase (GLUSy), which is a newly added reaction by the D3 expansion. PPCK is connected via polyenolpyruvate to 3-phosphoshikimate 1-carboxyvinyltransferase (PSCVT), another add by the D3 expansion. Furthermore, SUCDi is topologically connected with the added reaction ubiquinone L-Lactate dehydrogenase (L-LACD2), as cofactors ubiquinone-8 and ubiquinol-8 partake in both reactions. Interestingly, periplasmic glucose dehydrogen (GLCDpp), where ubiquinone-8 and ubiquinol-8 also participate, has a high DI as well. GLCDpp possibly causes its neighbouring reactions gluconokinase (GNK) and D-gluconate transport (GLCNt2rpp) to have high DIs too, due to stoichiometric coupling. These observations suggest that alterations in flux split ratios around important branching points - caused by network expansion - could result into higher DIs in reactions at their vicinities.

**Figure 6.**
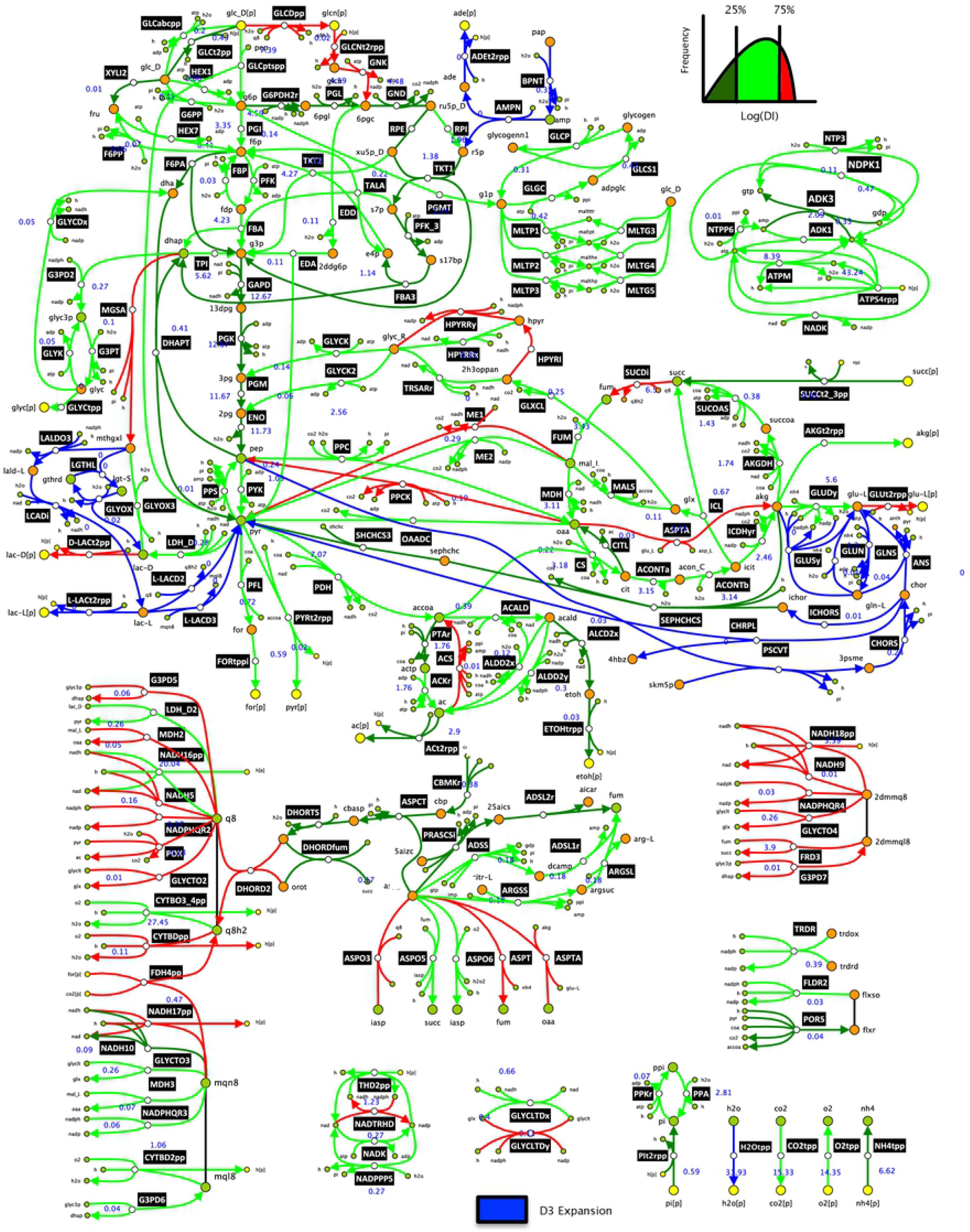
*E.coli* network diagram illustrating the deviation index (DI) of reactions when scaling up from D2 to D3. The flux values of the D3 flux profile (reference state) are given for the reactions in mmol/gDW/h next to their labels. The reactions added by the modular redGEM expansion from D2 to D3 are given in blue and the 307 enzymatic reactions in common between D2 and D3 are color-coded based on the DI. The reactions were categorized into low DI (0-25 percentile), medium DI (25-75 percentile) and high DI (75-100 percentile), and are given in dark green, light green and red, respectively. Diagram does not include all the reactions of the systems.

Overall, lower DIs were observed for reactions having a higher flux, pertaining to the core central carbon metabolism around which the models D1-3 were reduced (Figures 5 and 6). Since the cores of the reduced models contain the 12 precursor metabolites for biomass, their control patterns were expected to be similar. Stephanopoulos and Vallino point out that metabolic pathways of organisms have evolved over time to resist flux alterations at branching points [28]. The control architecture of an organism is built such that it preserves the flux splitting ratios of essential metabolic nodes. However, if two models have differences in the number of reactions and/or in the flux splitting ratios around an important branching point, the control architecture of the two systems can differ considerably.

Since we studied optimally grown *E.coli*, it was expected that the D1 to D2 expansion with the addition of XYLI2, F6PP and HEX7 would have influence on control patterns: the flux splitting ratios around the essential biomass precursor D-fructose 6-phosphate is altered. D-fructose 6-phosphate is a critical metabolic node for producing cell wall biomass building blocks and is located relatively upstream in the process of glucose catabolism. Altering flux splitting ratios around D-fructose 6-phosphate will have direct implications on the fate of the carbon flow across the whole network, particularly due to its upstream location.

The expansion from D2 to D3 results in different flux splitting ratios around three biomass precursors: pyruvate, polyenolpyruvate and 2-oxoglutarate. Again, we can expect flux control patterns across the models to differ as the proportion of carbon flow directed towards certain biomass building blocks is affected. However, within the central carbon metabolism, these precursors are located relatively downstream to the glucose uptake, compared to for instance D-fructose 6-phosphate. Consequently, we can expect that these flux splitting ratios have less impact on the growth control of the system than D-fructose 6-phosphate. If we were discussing the production of certain amino acids of interest, rather than just cellular growth, these ratios could be of higher importance to the analysis. The significance of a metabolic node is strongly subject to the scope of the study. Hence, it is difficult to imagine a “one-size-fits-all” model due to the complexity of the problems encountered in metabolic engineering.

Indeed, the importance of a metabolic branching point is very study-specific as objectives can vary significantly. Had we, for instance, been interested in the study of D-lactate production, it would have been essential to include the metabolism of methylglyoxal, D-lactate and L-lactate into the subsystems around which model reduction is performed. However, as we are not interested in the production of D-lactate, we are not that concerned about the high DI of D-lactate transporter (D-LACt2pp) when comparing D2 and D3 (Figure 6). Furthermore, if we were interested to produce D-lactate, it would be essential to consider implication of attempting to deviate flux towards the metabolism of D-lactate. If the redirection of flux towards D-lactate imposes important changes in the flux splitting ratios of significant metabolic nodes of wild-type *E.coli*, it may be worth considering other organisms that cause fewer modifications in flux distribution [28, 29].

#### Study of uncertainty in MCA

The MCA results of D1-3 were further studied by comparing their absolute deviations in the FCCs. We considered with respect to the central carbon subsystems to find which central carbon enzymes contributed in most uncertainty across the networks. The FCCs of reactions in the glycolysis (Figure 7) appear to have most absolute deviation stemming from enzymes in the glycolysis and in the PPP. When comparing D1 with D2 (Figure 7A), enzymes in glycolysis contribute most to the uncertainty whereas in the comparison of D2 and D3 (Figure 7B), the PPP contributes the most. Again, the additional connections around D-fructose 6-phosphate (Figures 1 and 2) when expanding from D1 to D2 could explain this. Differences in flux splitting ratio around D-fructose 6-phosphate affect the redistribution of the flux in the network and hence the control pattern. Generally, reactions with a larger flux exhibit less absolute deviations in their FCCs. This parallels the observation that central carbon reactions carrying higher flux are perhaps more rigid in control patterns.

**Figure 7.**
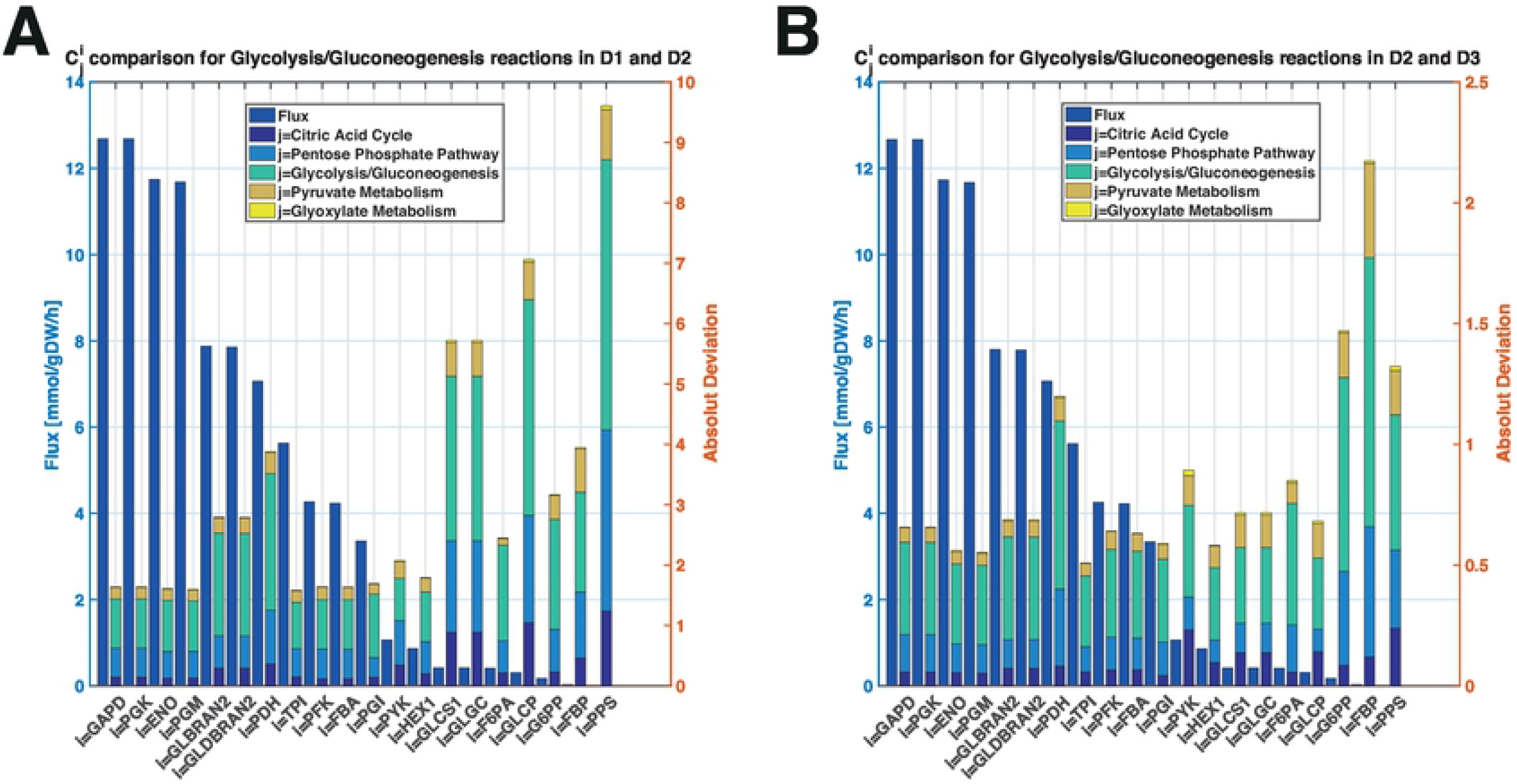
Comparison of flux and absolute deviations in FCCs for glycolytic reactions for (A) D1 versus D2, and (B) D2 versus D3. The absolute deviations computed per subsystem correspond to the sum of the absolute deviations in FCCs of reaction i with respect to all enzymes of the subsystem j. The reactions contained in a subsystem are as defined in the original GEM that was reduced [15]. The flux values shown did not deviate by more than 1% between pairs of models.

We perform a parallel analysis on FCCs of PPP reactions and similar observations were made. The expansion from D1 to D2 has most absolute deviations coming from enzymes from glycolysis (Figure 8A), whereas, the expansion from D2 to D3 has most deviations due to PPP enzymes (Figure 8B). Again, reactions carrying higher flux have less absolute deviation in their FCCs between the pairs of models. We analyzed FCCs individually in terms of absolute deviation (S5 Table), for both pairs D1 and D2 as well as, D2 and D3. PGI, TPI and PFK were the top three central carbon enzymes that resulted in the most absolute difference in flux control across the network. From the PPP enzymes, RPI resulted in the most absolute deviation in flux control. We also recall that RPI had sign-wise opposing control on cellular growth in the comparison of D1 and D2 (Figure 4A). Due to the highly non-linear nature of the studied systems, it is difficult to make direct conclusions on the causality of the observed deviations in control patterns of the networks. Most of the deviations were observed amongst peripheral transport reactions, rather than central carbon metabolism (S5 Table). Nevertheless, we could still find metabolic engineering decisions relevant to our study, independent of the complexity based on MCA outputs (Figure 4).

**Figure 8.**
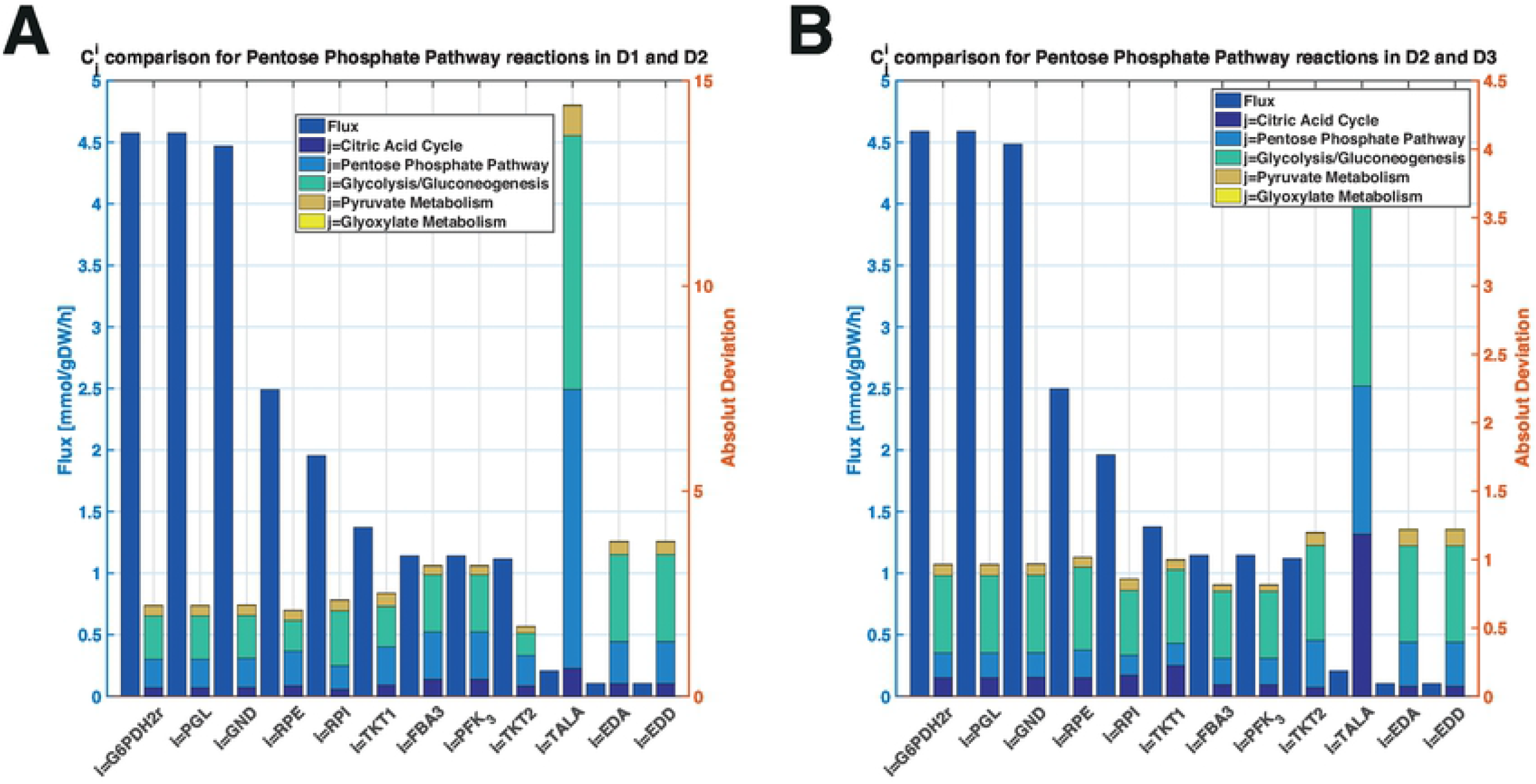
Comparison of flux and absolute deviations in FCCs for PPP reactions for (A) D1 versus D2, and (B) D2 versus D3. The absolute deviations computed per subsystem correspond to the sum of the absolute deviations in FCCs of reaction i with respect to all enzymes of subsystem j. The reactions contained in a subsystem are as defined in the original GEM that was reduced [15]. The flux values shown did not deviate by more than 1% between pairs of models.

## Conclusions

This work studied the impact of model complexity on the metabolic engineering decisions derived from MCA outputs. The redGEM and the lumpGEM algorithms were used to consistently reduce the *E.coli* iJO1366 GEM. Omics data for the physiology of optimally grown *E.coli* was integrated into the reduced stoichiometric models. The modularity of the reduced models assisted us in the development of a workflow allowing to preserve maximum equivalence between the flux profile and metabolite concentration steady-states. The ORACLE framework was used to generate populations of stable kinetic models around these reduced stoichiometric models. Our workflow ensured that we preserve equivalency amongst the populations of the kinetic parameters for the stable kinetic models. The MSCs were computed within the ORACLE framework for the populations of stable kinetic models. Analysis of the MSCs, revealed that we can derive context-specific metabolic engineering conclusions that are independent of the model’s complexity, as long as the reduction is performed consistently.

The “usefulness” of a kinetic model is highly dependent on the objectives of the study being undertaken. We selected the subsystems for the GEM reduction such that we: (1) cover the essential biomass precursor metabolites according to Neidhart as we focused primarily on cellular growth control and, (2) that we capture the ETC essential to account for redox potentials. The addition of reactions around D-fructose 6-phosphate when expanding from D1 to D2 appeared to significantly affect growth control patterns (Figure 4A). However, the expansion from D2 to D3 had considerably less impact as top cellular growth FCCs are consistent (Figure 4B). As D-fructose 6-phosphate is an essential precursor for cell wall fabrication, a network expansion affecting flux distribution around it can be expected to have significant impact on cellular growth control structure. Hence, it is essential to consider the importance of certain metabolic nodes with respect to the study goals in order to ensure no information is lost in the reduction. Again, importance of a metabolic node is strongly influenced by the nature and the objectives of the analysis.

The MCA summation theorem was used to postulate a deviation index (DI) that gave a numerical indication on the consistency of the FCCs with respect to a reaction. Most of the reactions around central carbon metabolism, carrying a higher carbon flux, appeared to have lower DIs. Flux control for reactions with larger fluxes were more robust, particularly if the number of connecting reactions did not change between models for the metabolites participating in the reaction. The larger DIs were noted in the ETC and peripheral reactions. Nevertheless, the consistency in the control patterns across reduced models allows us to make conclusions that are independent of the network complexity. In fact, we may not need full genome-scale kinetic models when the model reduction is done consistently as the essential, study-specific, information is accounted for by the reduced model.

This study demonstrates that systematic and modular model reduction algorithms ease the scale-up of kinetic models of metabolism. Our workflow describes an MILP formulation for insuring maximum equivalence between models when transferring steady-states. Furthermore, the workflow ensures that the kinetic parameters are kept as similar as possible between the populations of stable kinetic models built around the reduced stoichiometric models. To our knowledge, this is the first publication to date demonstrating transferability of steady-states between large-scale kinetic models, whilst obtaining consistent, study-specific, metabolic engineering decisions. As systematic model reduction algorithms gain momentum in the field, we hope to pave a path towards building more robust and transferable kinetic models for the community.

## Methods

We developed a workflow for building consistently reduced kinetic models from a genome-scale metabolic model (Figure 9). We used the redGEM algorithm to construct core models of increasing network size from the *E.coli* iJO1366 genome-scale model. The lumpGEM algorithm was used to generate lumped reactions for the biosynthesis of biomass building blocks (BBBs) for these models. We used thermodynamic-based variability analysis (TVA) [22] to study the flexibility of the models. We proposed a procedure for scaling up the flux and concentration steady-states from one model to another one using the MILP formulation. The ORACLE framework was enhanced, allowing us to keep parametric equivalency between the populations of kinetic models around the steady-states of the reduced models. These steps are further detailed below.

**Fig 9.**
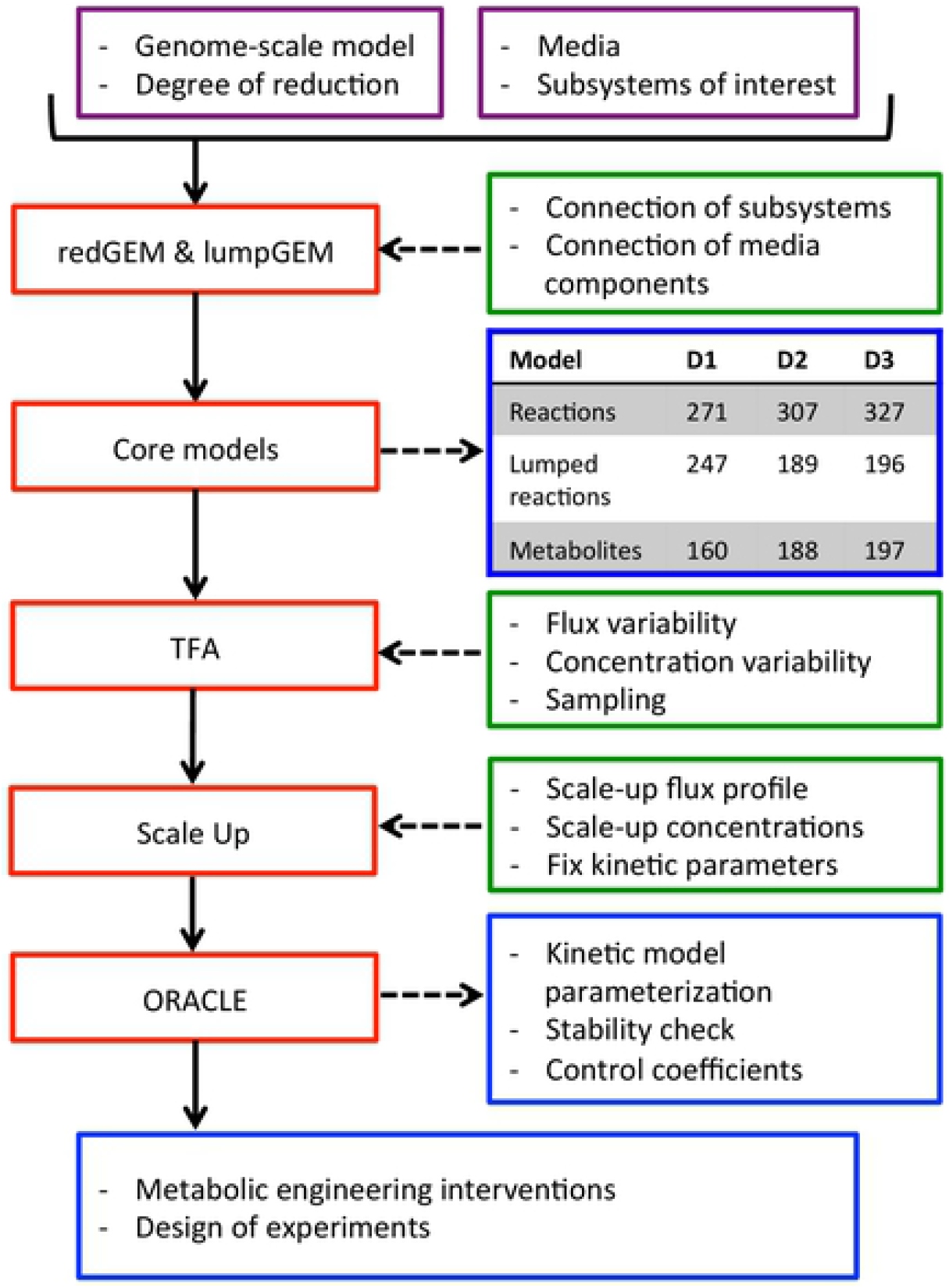
Workflow for building consistent reduced kinetic models. The various steps are discussed with further detail within the manuscript.

### Model reduction

The stoichiometry of the core networks was defined with the redGEM algorithm, which reduces systematically genome-scale model reconstructions of metabolism [7]. The *E. coli* iJO1366 genome-scale model was reduced, with aerobic minimal media, glucose as the sole carbon source, and the selected starting subsystems corresponding to central carbon metabolism (glycolysis/gluconeogenesis, citric acid cycle, pentose phosphate pathway, pyruvate metabolism, and glyoxylate metabolism). We incorporated all the reactions that use metabolites of the quinone/quinol pools (ubiquinone, ubiquinol, menaquinone, menaquinol, 2-dimethyl menaquinone and 2-dimethyl menaquinol) as the electron transport chain subsystem in order to account for the energy metabolism of the system. redGEM allows the user to define a degree of connection, D, to define the level of connectivity of the core. D is an input parameter of the redGEM and lumpGEM. D corresponds to the number of reaction required to connect the pairs of metabolites between starting subsystems, as defined in [7]. We generated core networks with a D of 1,2 and 3, which gave rise to models D1, D2 and D3 respectively. The lumpGEM algorithm [12] was used to generate lumped reactions for the biosynthesis of the BBBs for these core networks. Lumped reactions are sub-networks of reactions composed of non-core reactions that can be used to produce a BBB. Alternative lumped reactions were kept for each of the BBBs. Reactions that could not carry flux were considered as blocked and were removed.

For some of intracellular metabolites, a corresponding transport reaction has not been biochemically characterized and does not appear in the *E. coli* iJO1366 and in our reduced model. However, these metabolites, unless they are highly polar or very large, are subject to passive diffusive transport through the cell membrane. Therefore, we explicitly added transport reactions for these metabolites that operate at least at basal level (10^-6 mmol/(gDW*h)).

### Flux directionality assumptions

We make the following directionality assumptions for several bi-directional reactions:

1. Fructose-biphosphate aldolase (FBA) that is part of mid-lower glycolysis is set towards catabolism [18].
2. The bi-directional transports of magnesium and phosphate are both set towards uptake [19, 20].
3. Acetate kinase (ACKr) and phospho-transacetylase (PTAr) are both set towards the acetate production, because acetate is one of the main by-products [14].
4. The succinyl-CoA synthetase (SUCOAS) is set towards the production of succinate [14].

The polyphosphate kinases (PPK2r, and PPKr) are set towards the polyphosphate polymerization [21].

### Thermodynamic analysis

The available fluxomics and metabolomics data for the optimal growth of *E.coli* under aerobic conditions and minimal media was integrated in our models. The MILP formulation of the thermodynamics-based flux analysis was used to implement these data into D1, D2 and D3. Since the models were used to build kinetic models, it was undesirable for reactions to be at thermodynamic equilibrium, which would result in them having equal backward and forward fluxes. We imposed MILP constraints to ensure that the thermodynamic displacement, Γ [23, 26, 30], is not at equilibrium. For reactions near equilibrium Γ ≈ 1.

### Maximum equivalency between steady-states

We sampled the flux space of D1 in order to characterize the solution space without violating physiological, thermodynamic and directionality constraints. The convexity of the solution space enabled us to efficiently sample using the Artificial-Centering Hit-and-Run sampler in the COBRA Toolbox [31, 32]. We sampled flux vectors and used Principal Component Analysis (PCA) [33] to select a mean reference state. Similarly, we sampled and selected the reference state with PCA for the concentration solution space of this selected flux profile.

In order to make the comparison of the models equitable, we wanted to maintain most similar steady-states between the models. For instance, for D2 we would like the flux vector to be the equal possible to the one from D1. Topological differences in the models make it impossible to have numerically exactly the same flux distribution in larger model for the same reactions. Hence, we take the representative flux from D1 and apply it with percentage relaxation with upper and lower bounds, F^ub^_rxn,i_ and F^lb^_rxn,i_ respectively, into D2. Consequently, we use an MILP formulation to minimize the number of violations of flux boundaries that we are trying to impose. For each intracellular reaction that is shared between the two models we create a binary variable z_rxn, i_ so that when it is equal to 1, the constraints that we impose become inactive. We add for each of these reactions the following constraints:

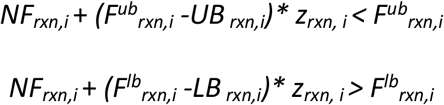

where, *UB* and *LB* are the upper and lower bounds of the net fluxes *NF* of the reactions. We minimize the sum of the binaries, *z_rxn, i_*, in order to have minimal violation of the flux constraints:

Minimize:

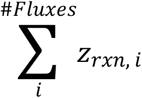

Subject to:

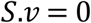

We implied a 1% relaxation to apply and test how many flux constraints we can impose without violation (minimal number of active binary variables *z_rxn, i_*). After applying the constraints that are not violating model boundaries of D2, we proceed to sampling the solution space. We selected a sample based on mean PCA as with the representative flux of D1. We then implied in a similar manner the concentration profile from D1 into D2 with a 1% relaxation and sampled the concentration space for this flux profile. We repeat this procedure when scaling up the flux and concentration steady-states from D2 into D3.

### Constructing kinetic models

We used the ORACLE framework to build 50’000 kinetic models around the steady-states for D1, D2 and D3. Available kinetic properties of enzymes from the literature [34] and the databases [35, 36] were incorporated. Reversible Hill kinetics [37] and convenience kinetics [38] were used for reactions with unknown kinetic mechanism (S6 Supporting information). Kinetic mechanisms with no or partial information about their parameter values were sampled within the space of kinetic parameters in the form of degree of saturation of enzyme [23]. We parameterized a population of kinetic models and performed consistency tests [23, 39, 40]. We then computed the flux and concentration control coefficients [23, 41]. For further details on the ORACLE workflow the reader is referred to [2, 3, 23–25, 30, 40, 42].

We preserved equivalency between populations of kinetic models for D1-3 by fixing the degree of saturation of enzymes from less complex models into the more complex models. We wanted to preserve model equality so that we can fairly compare MCA outputs of the models. Within the ORACLE framework, we added a feature for fixing the degree of saturation of enzymes. For the parameters that were common between D1 and D2, we fixed the degrees of enzyme saturations from D1 models into D2 models and we sampled the rest of the D2-specific parameters uniformly, until we found a stable model. Hence, we preserved equivalency of the kinetic parameters between D1 and D2. Analogously, we repeated this procedure to imply the degrees of enzymes saturations from D2 into D3.

### Control coefficient deviation index

In MCA the FCCs conform with the summation theorem defined in [26, 27]. The theorem implies that all the metabolic fluxes are systemic properties of the model and that their control is shared by all the reactions within the system. The summation theorem makes the assumptions that: (1) the parameters for which we compute flux control coefficients are of first order with respect to the flux, and that (2) the sum of a flux’s control coefficients with respect to all the parameters of the system is equal to one. We proposed a deviation index (DI) derived from the summation theorem to quantify the discrepancies in control patterns of a flux between two different models (Figure 10).

**Fig 10.**
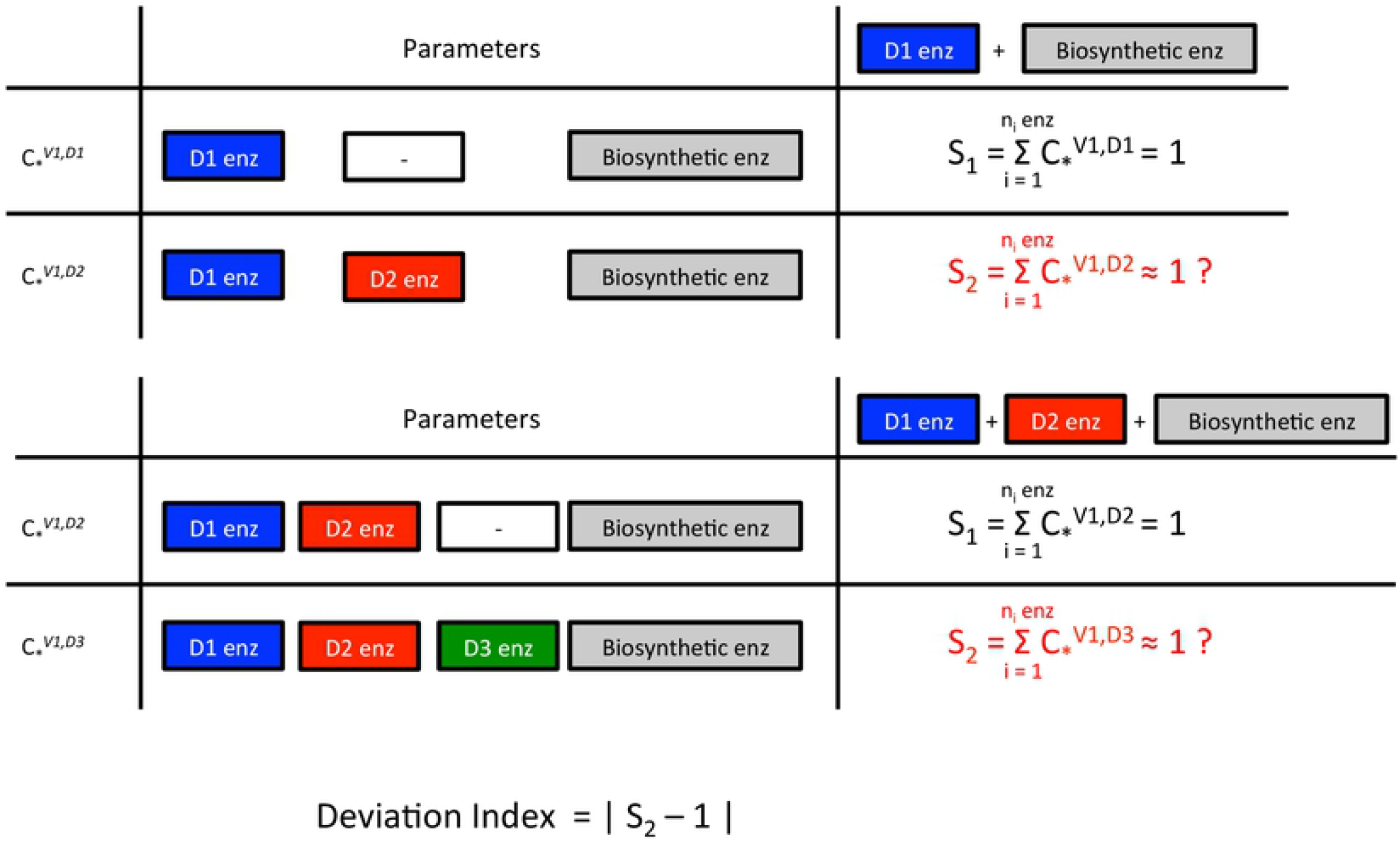
Derivation of the deviation index (DI) from the summation theorem.

## Supplementary

**S1 Table. Fluxomics and metabolomics data incorporated in the model.** Table with the fluxomics data in mmol/gDW/h and the concentration data in log(M).

**S2 Table. Thermodynamics-based variability analysis of models**. Spreadsheet with the list of metabolites and reactions inside the models with variability analysis of metabolic flux and metabolite concentrations.

**S3 Table. Flux and concentration steady-states**. Spreadsheet providing metabolic flux and concentration reference steady-states across the models with a comparative study.

**S4 Figure. Flux control coefficients of (a) glucose uptake, (b) formate excretion and (c) acetate excretion across the models**. Pairwise illustration of the union of the top 7 enzymes across the models in terms of absolute control over cellular growth for D1 versus D2, and D2 versus D3. The whiskers give the upper and lower quartiles of the FCC populations and the bars give the means.

**S5 Table. Analysis of absolute deviations in means of flux control coefficients for the entire systems**. Further assessment of deviations in flux control coefficients between model expansions from D1 to D2 and from D2 to D3.

**S6 Supporting information. Kinetic mechanisms used for the models.**

